# Tomographic Volumetric Bioprinting of Heterocellular Bone-like Tissues in Seconds

**DOI:** 10.1101/2021.11.14.468504

**Authors:** Jenny Gehlen, Wanwan Qiu, Gian Nutal Schädli, Ralph Müller, Xiao-Hua Qin

## Abstract

Tomographic volumetric bioprinting (VBP) has recently emerged as a powerful tool for rapid solidification of cell-laden hydrogel constructs within seconds. However, its practical applications in tissue engineering requires a detailed understanding of how different printing parameters (concentration of resins, laser dose) affect cell activity and tissue formation. Herein, we explore a new application of VBP in bone tissue engineering by merging a soft gelatin methacryloyl (GelMA) bioresin (<5 kPa) with 3D endothelial co-culture to generate heterocellular bone-like constructs with enhanced functionality. To this, a series of bioresins with varying concentrations of GelMA and lithium Phenyl(2,4,6-trimethylbenzoyl)phosphinate (LAP) photoinitiator were formulated and characterized in terms of photo-reactivity, printability and cell-compatibility. A bioresin with 5% GelMA and 0.05% LAP was identified as the optimal formulation for VBP of complex perfusable constructs within 30 s at high cell viability (>90%). The fidelity was validated by micro-computed tomography and confocal microscopy. Compared to 10% GelMA, this bioresin provided a softer and more permissive environment for osteogenic differentiation of human mesenchymal stem cells (hMSCs). The expression of osteoblastic markers (collagen-I, ALP, osteocalcin) and osteocytic markers (podoplanin, Dmp1) was monitored for 42 days. After 21 days, early osteocytic markers were significantly increased in 3D co-cultures with human umbilical vein endothelial cells (HUVECs). Additionally, we demonstrate VBP of a perfusable, pre-vascularized model where HUVECs self-organized into an endothelium-lined channel. Altogether, this work leverages the benefits of VBP and 3D co-culture, offering a promising platform for fast scaled biofabrication of 3D bone-like tissues with unprecedented functionality.

**Graphical abstract:** 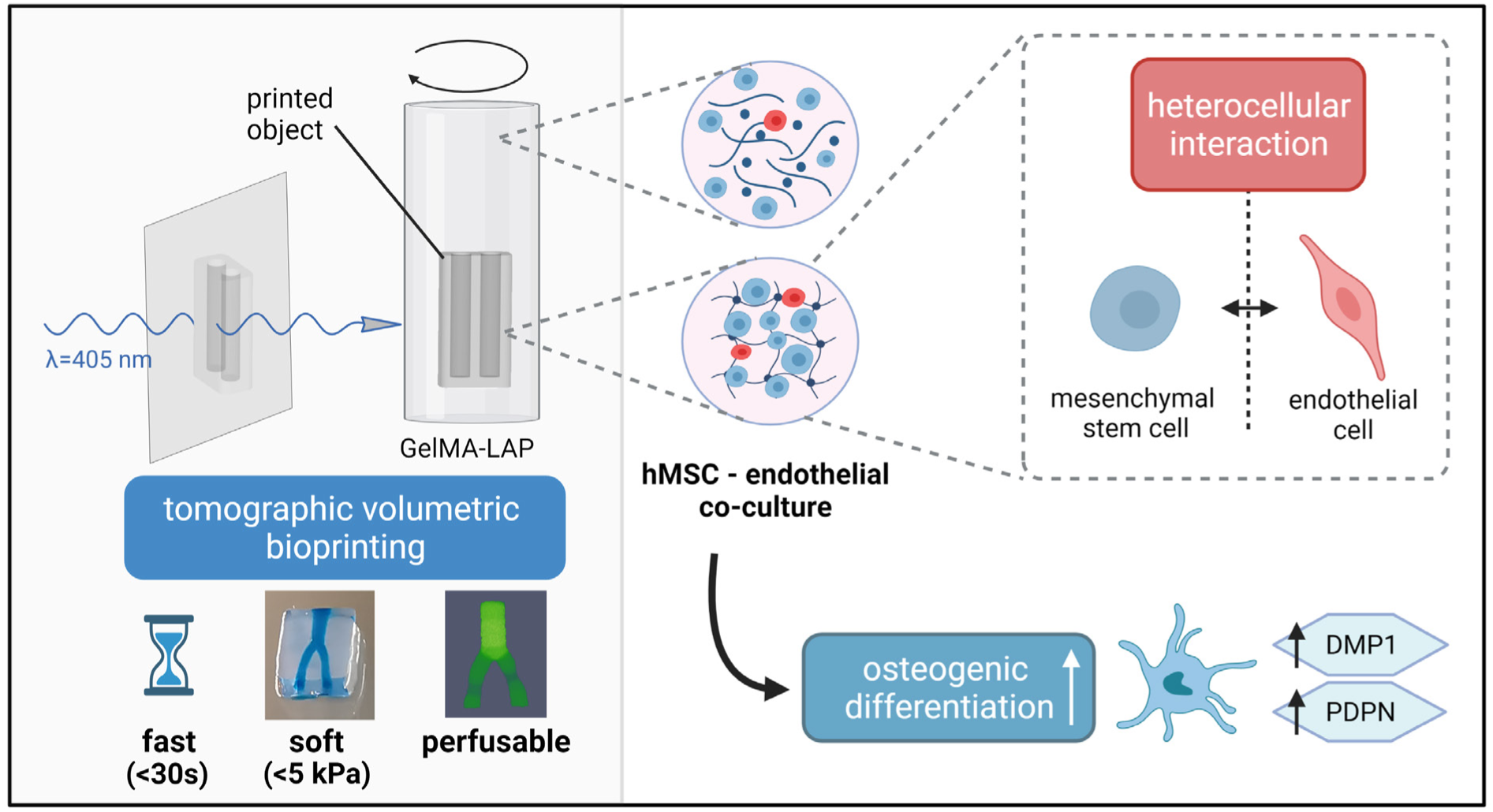

**Statement of significance:** This study explores new strategies for ultrafast bio-manufacturing of bone tissue models by leveraging the advantages of tomographic volumetric bioprinting (VBP) and endothelial co-culture. After screening the properties of a series of photocurable gelatin methacryloyl (GelMA) bioresins, a formulation with 5% GelMA was identified with optimal printability and permissiveness for osteogenic differentiation of human mesenchymal stem cells (hMSC). We then established 3D endothelial co-cultures to test if the heterocellular interactions may enhance the osteogenic differentiation in the printed environments. This hypothesis was evidenced by increased gene expression of early osteocytic markers in 3D co-cultures after 21 days. Finally, VBP of a perfusable cell-laden tissue construct is demonstrated for future applications in vascularized tissue engineering.

## 1. Introduction

The field of tissue engineering and regenerative medicine (TERM) aims to develop three-dimensional (3D) cellular constructs that mimic physiological tissue structure and composition [1, 2]. The idea of growing functional tissues *in vitro* holds the promise of revolutionizing future medicine by providing ‘spare parts’ for potential tissue replacements and drug screening for personalized treatments. 3D printing in combination with extracellular matrix (ECM)-mimicking materials has fueled recent advances in tissue engineering of complex biomimetic niches and has emerged as a promising biomanufacturing technology [3, 4]. Even though 3D bioprinting techniques based on layer-by-layer deposition have extensively evolved during the last decade, printing duration, scaffold porosity and clinically relevant construct size remain major challenges [5, 6].

Volumetric bioprinting (VBP) is a revolutionary technique based on tomographic light projections, for the fabrication of centimeter-scale constructs within tens of seconds [6-8]. This nozzle-free method leverages existing volumetric image modalities, such as computed tomography, to fabricate whole objects instantly and simultaneously. 3D light doses based on multi-angular object projections cumulatively induce crosslinking of photoresponsive cell-laden bioresins with high spatial precision. The advantages of VBP and its exceptionally high cell viability are especially relevant for applications in scalable biofabrication of living tissues. However, permissiveness and biocompatibility of employed hydrogel-based bioresins need to match the pace of cell differentiation and self-organization to eventually form a functional heterocellular tissue construct.

Stem cell-based tissue engineering has evolved rapidly and at present, osteogenic stimulants that drive the differentiation of mesenchymal stem cells (MSC) into osteoblasts are known [9-12]. *In vivo* differentiation is accompanied by matrix remodeling of the so-called osteoid, a hallmark of osteogenesis, starting with collagen deposition, alkaline phosphatase secretion and finally osteocalcin expression [13, 14]. The expression of the osteoblastic markers Runt-related transcription factor 2 (RUNX2), collagen type I (COL1a2), alkaline phosphatase (ALPL), matrix metallopeptidase 14 (MMP14) and osteocalcin (BGLAP), as well as the osteocytic markers dentin matrix acidic phosphoprotein 1 (DMP1), podoplanin (PDPN) and sclerostin (SOST) is a hallmark in the dynamic osteoblast-to-osteocyte transition. The terminal osteocytic stage and the formation of networks with a complex three-dimensional lacuna-canalicular architecture remains a major challenge for state-of-the-art *in vitro* models [15]. Recent studies report the optimization of biophysical parameters and the use of different scaffold materials to promote bone formation, but so far, the most advanced 3D human bone model only resembles early osteogenesis with low level expression of osteocytic marker genes [9, 16-18].

New concepts have to be explored to further enhance existing models of *in vitro* osteogenesis, one of them being a co-culture approach with endothelial cells. The lack of intercellular differentiation cues appeared as a major problem of *in vitro* cultures and previous studies have shown that the co-cultivation with macrophages and osteoclasts can enhance osteogenesis [19, 20]. Although the exact intercellular interactions remain poorly understood, a stimulating role of vascular endothelial growth factors (VEGF) in osteogenesis has been suggested [21]. VEGF is a signaling protein secreted by many cells that promotes the osteogenic differentiation of MSCs [22]. Additionally, other studies have shown the enhanced osteogenesis through endothelial co-cultures is associated with Wnt and bone morphogenic protein (BMP) signaling pathways [23]. Specifically, human umbilical vein endothelial cells (HUVEC) in combination with MSCs in the osteogenic lineage have been described as enhancers of osteoblast differentiation and function, based on their important role in the cellular communication network within the microenvironment of bone tissue [24, 25]. However, no long-term studies investigating the effect on terminal osteocytic differentiation have been conducted yet.

In this work, we report a new method to fabricate heterocellular bone-like tissues in seconds by leveraging the advantages of ultrafast tomographic VBP technique and 3D hMSC/HUVEC co-culture (**Figure 1**). Until now, only a handful of publications [8, 26, 27] reported VBP of cell-laden hydrogel constructs. These prior arts highlight that a VBP process is much faster than traditional layer-by-layer printing techniques such as extrusion bioprinting [16, 28]. This study sought to extend the applications of VBP for bone tissue engineering. First, a series of GelMA bioresin compositions were characterized with a focus on printability, mechanical properties, and cell-compatibility. Using a soft GelMA (<5 kPa), we provide an extensive functional assessment of volumetrically bioprinted heterocellular constructs by state-of-the-art cellular and molecular assays. The effects of intercellular interactions on osteogenic differentiation were examined in a longitudinal analysis in both 2D and 3D over 42 days. Finally, the VBP technique was employed to create a perfusable *in vitro* model where hMSC-containing constructs with imprinted channel structures were lined with endothelial cells for future developments of pre-vascularized bone tissues.

**Figure 1:**
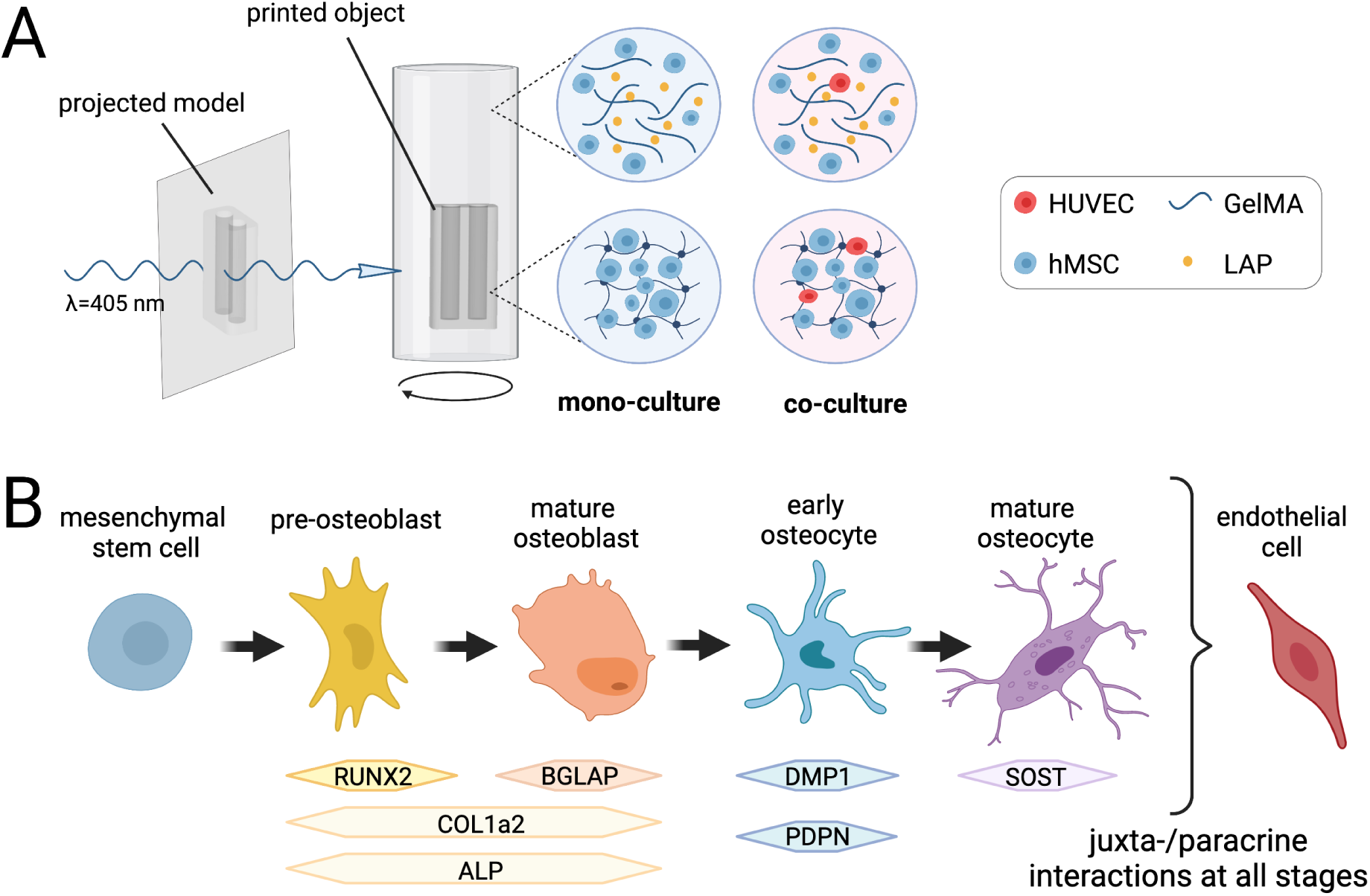
Illustration of tomographic VBP and enhanced stem cell osteogenic differentiation by endothelial co-culture. **A)** Schematic of VBP: Tomographic light projections lead to localized solidification of hydrogel constructs in a glass vial containing a cell suspension in a photo-crosslinkable bioresin (GelMA/LAP). Mixed cell suspensions allow the fabrication of self-organized heterocellular constructs combining human mesenchymal stem cells (hMSC) with endothelial cells (HUVEC). **B)** Schematic of osteogenic differentiation from MSCs to osteocytes with respective marker gene expression. The juxta- and paracrine interactions between all cells in the osteogenic lineage and endothelial cells promote osteogenic differentiation. Created with BioRender.com.

## 2. Materials and Methods

### 2.1 Biomaterials and volumetric bioprinting

#### 2.1.1 Bioresin preparation

All 3D constructs are based on gelatin methacryloyl (GelMA) which was synthesized following previously described protocols [29]. The degree of substitution (DS) of GelMA was estimated with ^1^H-NMR (Bruker Ultrashield, 400 MHz) in D_2_O. Integration signal (2.87–3.00 ppm) in GelMA was compared to unmodified gelatin lysine integration signal (2.87–3.00 ppm). Phenylalanine signal (7.1–7.4 ppm) was used as internal reference. DS was found to be ≈57% using the following equation:

DS = [1-(lysine methylene proton of GelMA)/(lysine methylene proton of gelatin)]*100%

To obtain semi-sterile materials for aseptic cell culture techniques, synthesized GelMA was sterile filtered and lyophilized through polytetrafluoroethylene (PTFE) membranes of tissue culture flasks with a pore size of 0.22 µm. For photoinitiated crosslinking of functionalized methacryloyl groups, GelMA was dissolved in PBS containing the photoinitiator Lithium-Phenyl-2,4,6-trimethylbenzoylphosphinat (LAP) in different concentrations ranging from 0.08 to 0.03% (w/v). To avoid loss of functionality dissolved GelMA was stored in the dark at 4°C.

#### 2.1.2 Volumetric bioprinting

For VBP, a printer prototype from Readily3D (EPFL Innovation Park, Lausanne, Switzerland) and the updated Tomolite v1.0 printer in combination with the updated Apparite software were used. To optimize printing settings and to achieve clearly defined constructs, laser dose tests were conducted for each GelMA resin. Defined spots were irradiated into a cuvette filled with solidified resin by the inbuild laser (λ=405 nm) for varying exposure times ranging from 2 to 64 seconds and varying average light intensities ranging from 1 to 32 mW/cm^2^. The light dose threshold (mJ/cm^2^) required for precise polymerization, but minimal off-target exposure, was calculated by multiplying exposure time with the average light intensity of the weakly visible polymerized spots in the cuvette. Light doses in the range of 100 to 600 mJ/cm^2^ were identified and used for different GelMA solutions depending on the resin batch, photocrosslinker concentration and cell number.

Construct printing was performed in sterile glass vials with 18 mm or 8 mm diameter in a volume of 3 ml or 1 ml resin, respectively. For cellular prints 3×10^6^ cells/ml resin were used. The required number of cells was pelleted by centrifugation after harvesting and directly immersed in prewarmed, liquefied (37°C) GelMA. After thorough mixing, the cell suspension was rapidly solidified at 4°C to prevent cell sedimentation and it was proceeded to printing immediately. The bioresin formulation remained solid during the whole printing process (< 60 s) although the bioprinter was stored at ambient temperature (22-23 °C). After printing, the cell suspension was warmed to 37 °C to liquefy the non-crosslinked resin, the printed construct was washed in warm PBS and transferred to medium for cultivation.

The predetermined refractive index of GelMA (1.37) was applied to acellular and cellular prints and a peak-to average power ratio (PAPR) of 6:1 was chosen to print with optimal resolution at reasonable printing speed. Construct models for volumetric printing were designed as Standard Triangle Language (STL) files using the stereolithography computer-aided design (CAD) software FreeCAD.

#### 2.1.3 in-situ Photo-Rheology

Rheological measurements of bioresins were performed with the rheometer MCR 302 (Anton Paar, Germany), using a parallel plate with 20 mm diameter. During time sweep measurements with an interval of 6 seconds for 5 minutes, UV-curing of GelMA resins was induced after 60 seconds by illumination with an UV-LED lamp (λ=365 nm, light intensity 35% (70% ≙ 20 mV/mm^2^), Thorlabs). The storage moduli (G’) and loss moduli (G”) were recorded to assess the crosslinking ability and terminal stiffness of the used resins. 45 µl of resin were loaded and the gap was set to 0.1 mm. To prevent drying, wet tissue paper was placed within the temperature chamber (25 °C). Measurements for each resin were triplicated.

#### 2.1.4 Scaffold mechanics

To characterize mechanical scaffold properties, the compressive modulus of cellular and acellular printed constructs was determined with a Zwick material testing machine (Zwick 1456, Germany). Unconfined uniaxial compression was tested with a preload of 5 mN, and a strain rate of 1 min^-1^ until a 50% maximal construct deformation was reached. Ramp compression at a speed ratio of 0.01 mm s^-1^ was applied to obtain stress-strain curves. Disk shaped triplicates (3-6 mm, h = 1-2 mm) were tested at room temperature and the compressive modulus was calculated within the strain range of 5% to 10%. Measurements of acellular samples was performed after incubation in PBS at 37°C for 24 hours and cellular samples were tested after cultivation in cell culture conditions on day 1, 7 and 28 after printing.

### 2.2 Cell cultivation

Cells were maintained at 37°C, 5% CO_2_, in T175 triple flasks and passaged at 80% confluency using 0.05% trypsin-EDTA solution (incubation for 5 minutes, 37 °C, gentle tapping to ensure detachment), followed by centrifugation at 300 g for 10 minutes at 4 °C. All cells were tested negative for mycoplasma contamination using a Universal Mycoplasma Detection Kit (ATCC).

#### Human Mesenchymal Stem Cells (hMSC)

hMSCs (Normal Human Bone Marrow Derived Mesenchymal Stem Cells, pooled donor, Lonza) were used for experiments in passage 4 to 7 and were cultivated in Dulbecco’s Modified Eagle’s Medium (DMEM, Gibco) containing 10% fetal bovine serum (FBS, Gibco) and 1% antibiotic-antimycotic solution (Penicillin-Streptomycin-Fungizone, Gibco). For cell expansion, medium was supplemented with 1% MEM non-essential amino acids (Gibco) and 0.001% basic fibroblast growth factor (bFGF, Gibco, Thermo Fischer Scientific). To induce osteogenic differentiation, medium supplemented with ascorbic acid (50 µg/ml, Sigma-Aldrich), dexamethasone (100 nM, Sigma-Aldrich) and beta-glycerophosphase (10 mM, Acros Organics, Thermo Fischer Scientific) was used.

#### Human Umbilical Vein Endothelial Cells (HUVEC)

HUVECs (Human Umbilical Vein Endothelial Cells, pooled donor, Lonza) were used in passage 2 to 6 and were cultivated in Endothelial Cell Growth Medium-2 (EBM-2 medium supplemented with EGM-2 SingleQuots supplements, Lonza).

#### Endothelial Co-culture

hMSCs and HUVECs were harvested separately and mixed in a 5:1 ratio. For VBP, cells were randomly mixed with the resins by careful pipetting. Fully supplemented osteogenic hMSC medium and EGM-2 medium was mixed in a ratio of 1:1 and used for cultivation of co-cultures.

### 2.3 Perfusable vascularization model

3D constructs with perfusable channels (Ø <1mm) were designed and printed in a hMSC-containing cell suspension (3×10^6^ cells/ml). After cultivation for 7 days in osteogenic medium, HUVECs (5×10^6^ cells/ml) were suspended in Collagen I gel (1 mg/ml, pH 7.4) and loaded into the perfusable channels using a blunt-tip cannula, (Ø 0.8 mm). To trace HUVECs in co-culture, cells were labelled with the live-cell labelling dye Vybrant DiD (Invitrogen). Therefore, 1ml cell suspension with 5×10^6^ cells/ml was mixed with 25 µl cell-labeling solution (1mM) and incubated for 30 minutes at 37°C. The capability of endothelial cells to self-organize was exploited to create an endothelial lining of the channels and constructs were imaged with a confocal microscope 7 days after seeding. Constructs were immersed for 30 minutes in the cell-permeant dye calcein-AM (2 µM in PBS) before seeding to label hMSCs.

### 2.4 Imaging and Image analysis

#### 2.4.1 Microscopy

2D cultures and 3D constructs were imaged using a confocal laser scanning microscope (Zeiss LSM880) and overview images of acellular constructs were taken with a stereomicroscope (Leica Stereo M205 FA). The respective software ZEN and LAS X were used. Other acellular construct images were captured with a macro lens camera (Xiaomi). When imaged in air, constructs were transferred to a dry surface and surface water was wiped with tissue paper.

Images for viability assessment were quantified using the automated particle analysis plugin on ImageJ (NIH). The 3D rendering software Imaris (Oxford Instruments) was used to visualize z-stacks as 3D-rendered maximum intensity projections.

#### 2.4.2 Staining of 3D constructs

To enhance contrast of 3D constructs, different dyes were used for staining. Gels were incubated for up to 24 hours until sufficiently stained in 10% alcian blue solution in PBS for brightfield imaging, or in 1% EosinY dissolved in PBS for fluorescent imaging.

#### 2.4.3 Microcomputed tomography (microCT)

Scans with a voxel resolution of 34 µm were performed in air with a micro-CT45 device (Scanco Medical) with the following parameters: energy = 45 kVp, current 17 µA, integration time 600 ms. The micro-computed tomography (micro-CT) images were Gaussian filtered (σ = 1.2) and then threshold using a grayscale value corresponding to −10 mg hydroxyapatite/cm^3^. The greatest connected component was chosen to remove structures not belonging to the printed construct. Since the hydrogel had a relative low attenuation property, the 3D image was further smoothened using each two iterations of binary dilation and erosion. Then, a rectangular volume of interest was generated that lies completely within the boundaries of the 3D image to analyze the channel thickness. The channel thickness was determined by filling maximal spheres into the channel using the distance transformation method [30]. These image processing steps were done using an in-house developed software framework based on Python 3.6.6 (Python Software Foundation) using the SciPy 1.2.0 library and IPL software (V5.42 Scanco Medical). The 3D renderings were generated with ParaView.

The ECM mineralization of cellular constructs with mono- and co-cultures (each n =4) were monitored by weekly micro-CT scans for 6 weeks using the same scanning settings. To avoid scaffold moving, samples were fixed between mesh holders and cultivated in bioreactors with a volume of 5 ml. The bioreactors were outside of the incubator during the scanning time of ca. 40 min. the reconstructed images were gaussian filtered (σ = 1.2) and the mineralized scaffold was distinguished from the background using a threshold of 114 mg hydroxyapatite/cm^3^.

### 2.5 Functional analysis of bioprinted constructs

#### 2.5.1 Cell Viability and Morphology

Cell viability was assessed by performing live/dead assays using the cell-permeant dye Calcein-AM (1 mM stock in DMSO, used 1:500 in PBS) and the cell-impermeant dye Ethidium homodimer-1 (2 mM stock in DMSO, used 1:1000 in PBS). 2D and 3D cultures were incubated for 10 or 30 minutes respectively at 37°C, subsequently washed with medium and imaged within one hour.

Cellular morphology was assessed at different timepoints after fixation with paraformaldehyde (PFA) (4%, 15/30 min for 2D/3D cultures respectively, RT). Unspecific binding sites were blocked for >1 hour with 1% bovine serum albumin (BSA) and cell membranes were permeabilized with 0.2% Triton-X100 in PBS (0.1% BSA) for 10 minutes. Cell nuclei and the actin cytoskeleton were stained with Hoechst (1 mg/ml, 1:1000) and Phalloidin tetramethyl rhodamine (TRITC) (1:1000) in PBS (0.1% BSA) (>2 hours, RT). Samples were kept dark and stored at 4°C in PBS.

#### 2.5.2 Quantification of gene expression

##### RNA isolation

Samples for gene expression quantification were collected after cultivation for 1, 7, 14, 21, 28, 35 and 42 days. RNA was directly isolated from 2D cultures (1×10^5^ cells/10cm^2^, n=3) following a TRIzol-based protocol. For cell lysis 1 ml TRIzol was added to each sample and incubated for 5 minutes at RT. The cell suspension was transferred into a reaction tube, mixed with 200 µl Chloroform and vortexed for 15 seconds. After 3 minutes samples were centrifuged at full speed (>12000 g) for 15 minutes and the upper water phase containing RNA was aspirated and transferred into a precooled tube. An equal volume of isopropanol was added, and samples were incubated for 10 min followed by 10 minutes centrifugation. The supernatant was discarded, the precipitate was washed with 80% ethanol (v/v) and centrifuged again for 5 minutes. The supernatant was removed, and samples were left open to dry completely for at least 10 minutes. The precipitated RNA was dissolved in 20 µl RNase-free water.

Printed 3D constructs for RNA analysis were segmented with a scalpel into smaller pieces immediately after printing (50 mm^3^ with 3×10^6^ cells/ml) and were subsequently treated as independent samples. RNA was isolated from 3D samples (n=2-3) by combining a TRIzol-based cell lysis with a RNeasy Mini Kit (Qiagen). Samples were collected at aforementioned timepoints, snap frozen in liquid nitrogen and stored at −80°C until further usage.

For RNA isolation 300 µl TRIzol were added to the frozen 3D samples and the gels were dissociated in 1.5 ml reaction tubes using a rotating pellet pestle grinder. Afterwards additional 400 µl of TRIzol were added, samples were incubated for 5 minutes at RT and centrifuged at full speed (>12000 g) for 5 minutes. To remove gel remnants that could block spin columns the supernatant was transferred to QiaShredder columns (Qiagen) and centrifuged for 2 minutes. 700 µl of 70% ethanol (v/v) was mixed with the flow through and the total volume was transferred to RNeasy spin columns. The following steps were performed according to manufacturer’s instructions including DNase I digestion. RNA was eluted in 20 µl RNase-free water. The RNA concentration of all samples was measured using a NanoDrop spectrophotometer (λ =260 nm) and samples were stored at −80°C until further usage.

##### cDNA synthesis and qPCR

500 ng or 200 ng of isolated RNA from 2D and 3D samples, respectively, were reverse transcribed into cDNA in a volume of 10 µl using the PrimeScript RT Master Mix (Perfect Real Time, TaKaRa) (37°C, 15 minutes; 85°C, 5 seconds).

20-25 ng cDNA was subsequently used for gene expression analysis via qPCR using TaqMan Fast Universal PCR Master Mix (2x), no AMP erase UNG (95°C, 20 seconds; 43 cycles of 95°C, 1 second; 60°C, 20 seconds) in a CFX96 real-time PCR system (BioRad). Expression levels of two housekeeping genes (β-actin (ACTB), glyceraldehyde 3-phosphate dehydrogenase (GAPDH)) were used to normalize expression levels of eight osteogenic marker genes (RUNX2, BGLAP, DMP1, ALPL, PDPN, SOST, COL1a2, MMP14) which were analyzed with specific TaqMan probes (Applied Biosystems, see **Table S1**).

Obtained data was analyzed for relative gene expression after normalization to two housekeeping genes(ΔCt) with the BioRad CFX Maestro software and the analysis software qBase+ (BioGazelle).

##### ALP/DNA quantification

Activity levels of alkaline phosphatase (ALP) in relation to DNA expression levels were quantified as marker for osteoblast differentiation after cultivation for 1, 7, 14, 21, 28, 35 and 42 days. ALP activity quantification in the crude lysate of 2D (1×10^5^ cells/10 cm^2^, n=3) and 3D (50 mm^3^ with 3×10^6^ cells/ml, n=2-3) samples was performed according to manufacturer instructions of the Alkaline Phosphatase Assay Kit (colorimetric) (abcam).

2D samples were collected at aforementioned timepoints by adding 100 µl of provided ALP assay buffer to washed 2D cultures and cells were harvested using a cell scraper. 3D samples were directly transferred into 100 µl ALP assay buffer. All samples were immediately snap frozen in liquid nitrogen and stored at −80°C until further usage. After thawing, samples were dissociated using a rotating pellet pestle grinder. The ALP assay was performed in duplicates for each sample and standard.

Samples used for the ALP assay were kept at RT for 48 hours, centrifuged for 10 mins (>12000 g) and subsequently, 12.5 µl of the supernatant was used for DNA quantification using Quant-iT PicoGreen dsDNA Quantification Kit (Invitrogen) following the manufacturer’s instructions. Standards containing 0, 10, 50, 250 and 1000 ng DNA/ml were prepared and measured in a total volume of 100 µl using a plate reader (480 nm/520 nm).

To correct for differences in cell number, ALP activity was normalized to the DNA content. For co-cultures, the contribution of HUVECs to the total DNA was subtracted based on the initial co-culture ratio of 5:1.

### 2.6 Statistical Analysis

Statistical data analysis was performed using GraphPad Prism 8.2.0. Normality was assumed due to low sample numbers and parametric tests were performed to assess significance. Student’s t-test or Analysis of Variance (ANOVA) were applied for pairwise comparison or global comparison between experimental groups, respectively. The Holm-Sidak method was applied to correct for multiple unpaired comparisons using t-tests which assume a similar data scattering among populations at different timepoints. Significance of p-values is defined as α < 0.05 (*), < 0.01 (**) and < 0.001 (***).

## 3. Results and Discussion

### 3.1 Bioresin optimization for volumetric bioprinting

This study explored the use of VBP for generating heterocellular 3D constructs that exhibit enhanced osteogenic differentiation *in vitro* (**Figure 1**). The ability of volumetric bioprinter from Readily3D [6] to rapidly fabricate 3D cell-laden constructs elucidated its superiority to traditional extrusion-based 3D printers. This supremacy is justified by the very short printing time (<1 min) for centimeter-scale constructs with micrometer-scale resolution (**Supplementary Video S1**).

The VBP process is illustrated in **Figure S1A**. After cell harvesting, hMSCs were mixed with GelMA/LAP resin formulations at 37 °C in a glass vial. Next, the bioresins were cooled down to 4 °C for 15 min to form a physically crosslinked cell-laden gel before bioprinting at ambient temperature. Since the bioprinting process was fast, the resin formulation remained solid. After warming to 37 °C for 10 min, the printed constructs were obtained and transferred to cell culture media.

Firstly, we sought to characterize and identify optimal bioresin compositions for hMSC-based bone tissue engineering that support a high level of cell viability, 3D osteogenic differentiation and structural stability of the hydrogels after bioprinting. The rationale for GelMA as bioresin of choice is based on its excellent cell-compatibility and its frequent use in 3D cell culture [31-35]. The printing of different centimeter-scale 3D models (**Figure 2, Figure S1B**) with reliable performance was achieved using resins with GelMA concentrations of 2.5%, 5% and 10% in combination with varying photoinitiator (LAP) concentrations (0.03%, 0.05%, 0.08%). Constructs with hollow channels (Ø ≤ 1 mm) (**Figure 2A, B**) were designed and printed with the aim to be adoptable for future perfusion dynamic culture studies. High-resolution 3D reconstructions of the printed models were generated through micro-CT scans with a voxel resolution of 34 µm, and were used to visually confirm the printing precision. The STL model channel diameters in the range of 1 mm could be precisely recreated, although scaled shrinkage of the printed constructs was observed after incubation in PBS (**Figure 2A**, iv = 1 mm, **B**, i,ii,iii = 1.2 mm before and 0.9 mm after bifurcation). These findings show that soft hydrogel constructs with hollow architecture, which traditionally could only be printed with additional supporting structures in multiple steps, can be easily fabricated in single step by VBP within seconds.

**Figure 2:**
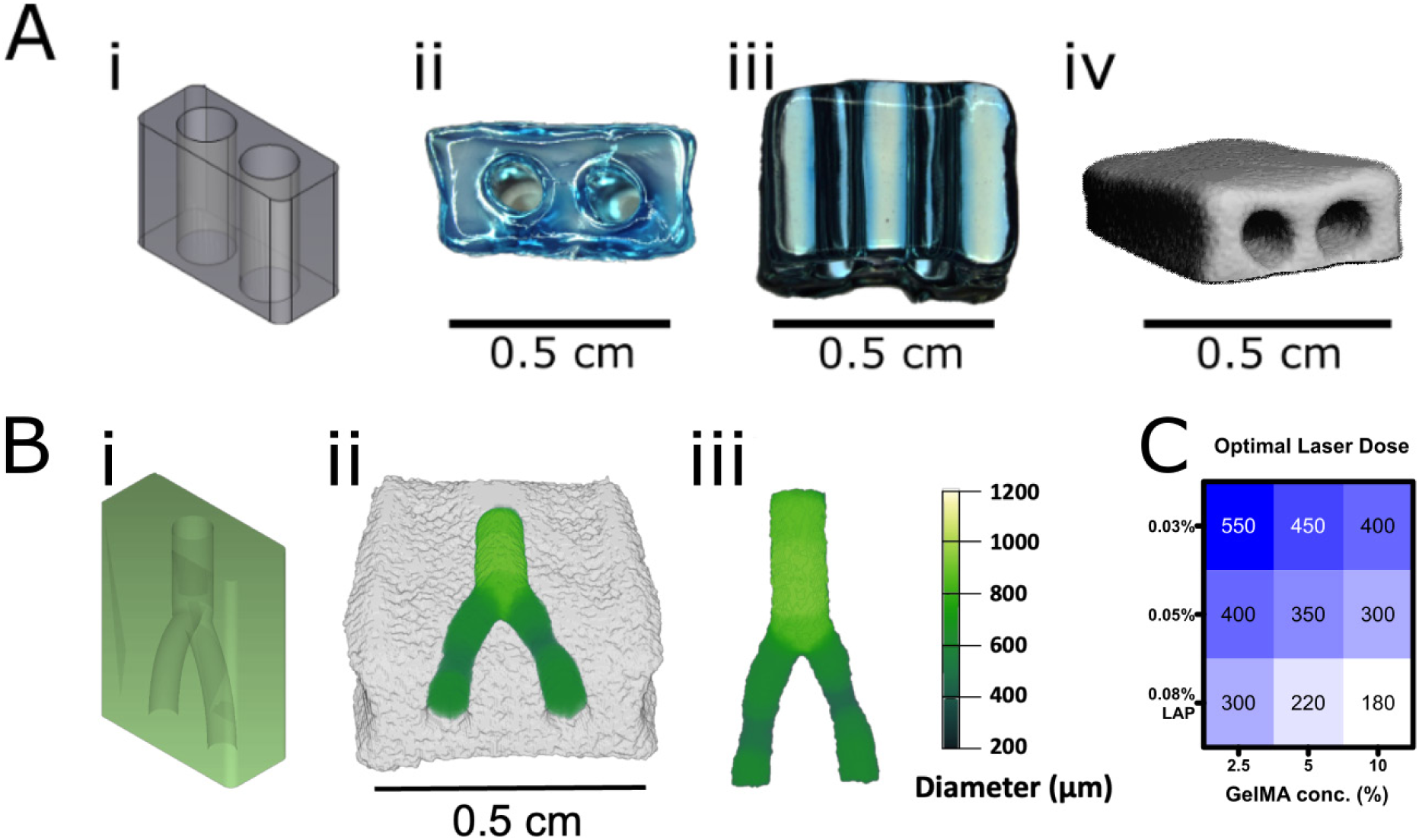
VBP of *in vitro* models with a hollow channel architecture. **A)** Volumetric printing of a simplified 3D model with hollow channels: the CAD model (i), top view (ii), side view (iii); and a 3D reconstruction by microCT scannning (iv). Printed samples were stained with Alzian blue to increase contrast. **B)** 3D rendering of a microCT scan of a branched channel model. CAD model (i), 3D volume reconstruction (ii), dissected channel (iii). Green color coding represents the channel diameter. A bioresin with 5% GelMA and 0.05% LAP was used for all gel constructs. **C)** Optimal laser doses for volumetric printing of resin formulations with different GelMA/LAP concentrations.

To achieve high print fidelity, the optimal laser settings have to be determined and validated regularly because the functionalization degree and reactivity of GelMA bioresins show batch variation, and changes of physicochemical properties within one batch with ongoing warming cycles and premature polymerization. To mimic the volumetric printing process, a multi-step rheology measurement of a resin (5% GelMA, 0.05% LAP) was performed (**Figure S1C**). By cooling the sample from 37 °C to 4 °C at a rate of 1 °C/min, both storage (G’) and loss modulus (G’’) was monitored. A sol-gel transition was observed at 16 °C. Subsequently, the sample was exposed to UV-LED 365 nm irradidation (10 mW/cm^-2^) for covalent crosslinking of GelMA via free radical polymerization, which resulted in a rapid increase in shear modulus (G’ ∼ 3.2 kPa, G’’ ∼ 450 Pa) after 5 min irradiation.

Laser dose tests spanning a broad range of average light intensities were used to identify an appropriate light dose that enables highest printing resolution. The required light dose also depends on the power of the inbuilt lasers in the different printer prototypes used in this study, as well as on the diameter of the printing vessel (8 mm/18 mm). Generally, required light doses are inversely correlated to GelMA and LAP concentration whereby the latter has a greater effect (**Figure 2D**). The required light dose for printing a bioresin with 10% GelMA and 0.08% LAP was approximately 180 mJ/cm^2^, whereas the dose required for another bioresin with 2.5% GelMA, 0.03% LAP was approximately 550 mJ/cm^2^. All other tested GelMA/LAP concentration combinations were printable within this range (**Figure 2D**). The required light dose directly correlates with the printing duration. However, the printing time does not increase when scaling up the construct volume as long as the same irradiation intensity is supplied [6]. Nevertheless, even light doses above 500 mJ/cm^2^ achieved complete polymerization of constructs with a volume of more than 10 cm^3^ in less than one minute.

Although a broad range of GelMA/LAP concentrations were printable, only a few resin compositions were suited for elaborate experimental procedure requiring manual handling. While the seminal study on VBP by Bernal et al. reported a stiff matrix (24 kPa) made of 10% GelMA, we reasoned that a lower mechanical stiffness and thus enhanced matrix permissiveness are essential for cell-matrix remodeling and hMSC differentiation [28, 36, 37]. Construct stability mainly depends on GelMA concentration. Bioresins containing 5% GelMA were chosen for future experiments as a good tradeoff between lower stiffness compared to 10% GelMA, and improved gel stability compared to 2.5% GelMA. Furthermore, the printability of a bioresin containing 2.5% GelMA and 2.5% sacrificial gelatin (total gelatin concentration of 5%) was confirmed. Aiming for an *in situ* softening environment, these constructs are expected to exhibit time-dependent decrease of gel stiffness and enlargement of porosity with ongoing incubation and resultant release of gelatin. Interestingly, all GelMA constructs with sacrificial gelatin at varying LAP concentrations showed enhanced ALP expression compared to the control group, implying the improved permissiveness. However, it is important to note that this improvement was accompanied with the compromise on surface smoothness (**Figure S2**). With this information in mind, 5% GelMA without sacrificial gelatin was determined as the optimal resin for further experiments since it combines high print fidelity and ease of handling. Additionally, cellular constructs with 5% GelMA showed enhanced osteogenic differentiation reflected by the increased relative gene expression of the osteocytic marker gene PDPN compared to constructs with a GelMA concentration of 10% (**Figure S3**).

### 3.2 Effect of photoinitiator concentration

We investigated whether the LAP concentration in combination with the respective light dose influences printing fidelity as indicated by surface smoothness and cell-compatibility. Construct staining with the freely diffusible, fluorescent dye Eosin Y allowed confocal microscopic imaging at high resolution (**Figure 3A**). From a side view, linear structures could be observed as printing artifacts at any concentration but no differences in surface smoothness from neither side (**Figure 3B**, i-iii) nor top view (**Figure 3B**, iv-vi) could be observed. This observation suggests that GelMA-based bioresins have excellent printability provided the light dose and photoinitiator concentrations are carefully selected.

**Figure 3:**
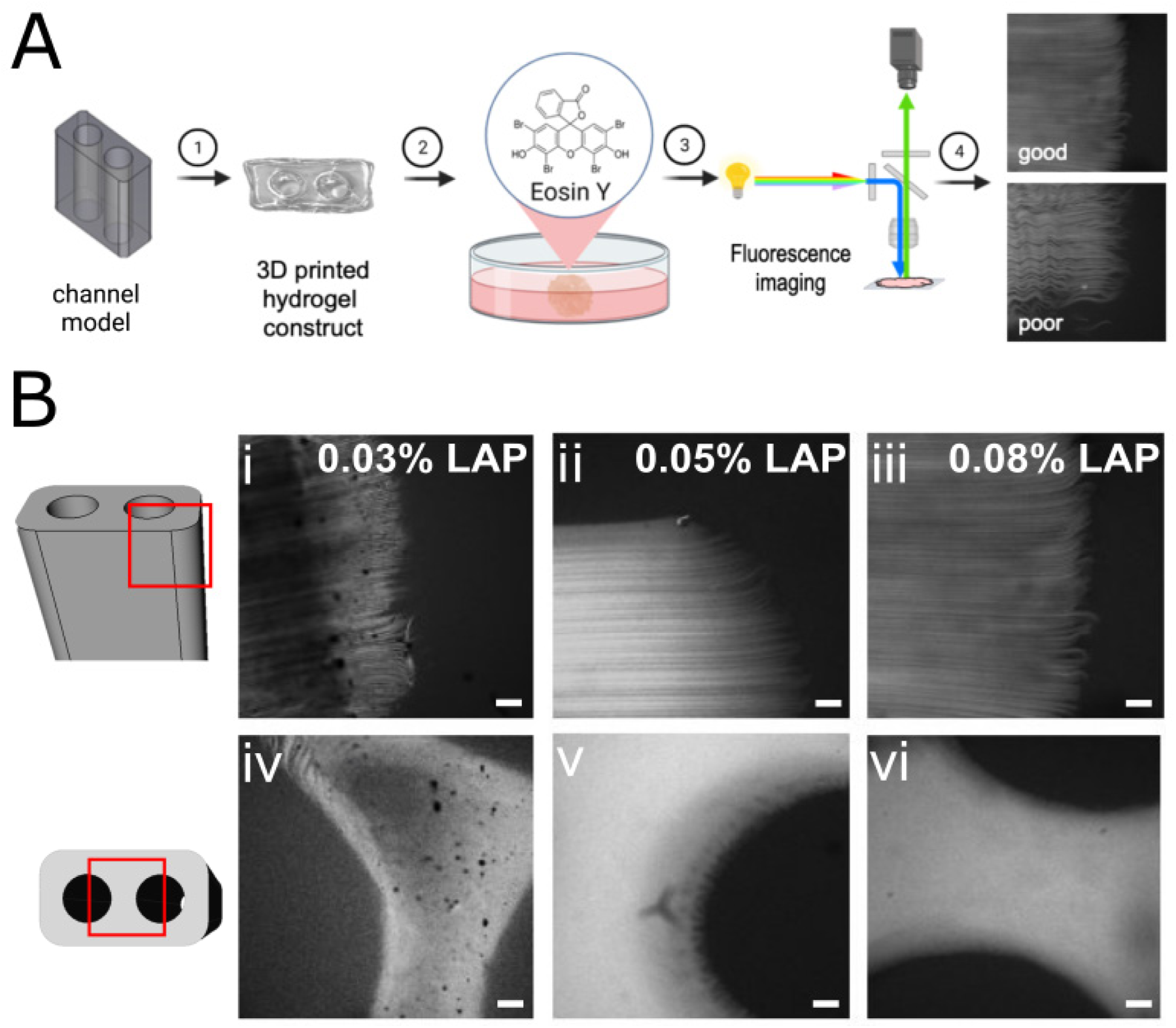
Assessment of the surface smoothness of printed constructs by Eosin Y staining and confocal fluorescence microscopy. **A)** Schematic workflow of the surface assessment in volumetrically printed 3D constructs. Printed constructs were incubated in EosinY solution for staining and then imaged with confocal laser scanning microscopy. Top and side view images revealed printed surfaces with good and poor quality (matrices in light grey). Created with BioRender.com. **B)** Confocal fluorescence microscopy images after Eosin Y staining of volumetrically printed constructs revealing the structural fidelity of printed constructs as a function of LAP concentration (side view: i, ii, iii; top view: iv, v, vi), scale bars = 100 µm.

Thereafter, we investigated the physical properties of hydrogel constructs by *in situ* photo-rheology and Zwick mechanical characterization to test if matrix stiffness affects cell spreading (**Figure 4 A-C**). The crosslinking efficiency of the bioresins were assessed by time-lapsed *in situ* photo-rheology with UV-365nm irradiation (**Figure 4A-B**). The evolution of storage (G’) and loss (G’’) moduli was monitored at an interval of 10 s. All compositions reached a G’-plateau in the range of 850 Pa – 1850 Pa (**Table 1**) after an irradiation for 4 min. No significant differences in the G’-plateau moduli were observed between the 0.03% and 0.05% LAP groups. However, the gels with 0.08% LAP exhibited the highest stiffness. Moreover, the sample with 0.03% LAP reached the onset (G’ > 5 Pa) at ca.116 s, whereas the onset time for the other two groups was significantly shorter (**Figure S5A**). Nevertheless, a comparison with t_90%-G’,_ the time required to reach 90% of the G’-plateau value, showed no significant difference (**Figure S5B**). Together, the 0.03% LAP group has the lowest photoreactivity, whereas higher LAP concentrations accelerated the photocrosslinking.

**Table 1.** Effect of LAP concentration on the photocrosslinking of GelMA. Results determined by *in situ* photo-rheology: 5% GelMA, UV-365 nm at 20 mW/cm^2^, 4 min.

| LAP (%) | G'-plateau<sup>a</sup><br>(Pa) | G''-plateau<sup>a</sup><br>(Pa) | loss factor<br>(x 10<sup>-3</sup>) | t<sub>onset</sub><br>(s) | t<sub>90%-G'</sub><br>(s) |
| --- | --- | --- | --- | --- | --- |
| **0.03%** | 1341.6 ± 16.2 | 7.4 ± 3.0 | 5.5 ± 2.3 | 116 ± 3.5 | 276 ± 0 |
| **0.05%** | 1155.8 ± 273.9 | 4.7 ± 3.4 | 3.9 ± 2.2 | 96 ± 10.4 | 264 ± 6.0 |
| **0.08%** | 1709.9 ± 137.8 | 18.1 ± 8.6 | 10.4 ± 4.1 | 90 ± 0 | 274 ± 13.9 |

<sup>a</sup> G', G'' and loss factor are the plateau values at 300 s;
<sup>b</sup> t<sub>onset</sub> is defined at the first data point where G'(t) > 5 Pa, indicating the time needed for curing.
<sup>c</sup> t<sub>90%-G'</sub> is defined at the time point where G'(t) reached 90% of the G' values at 300 s.

**Figure 4:**
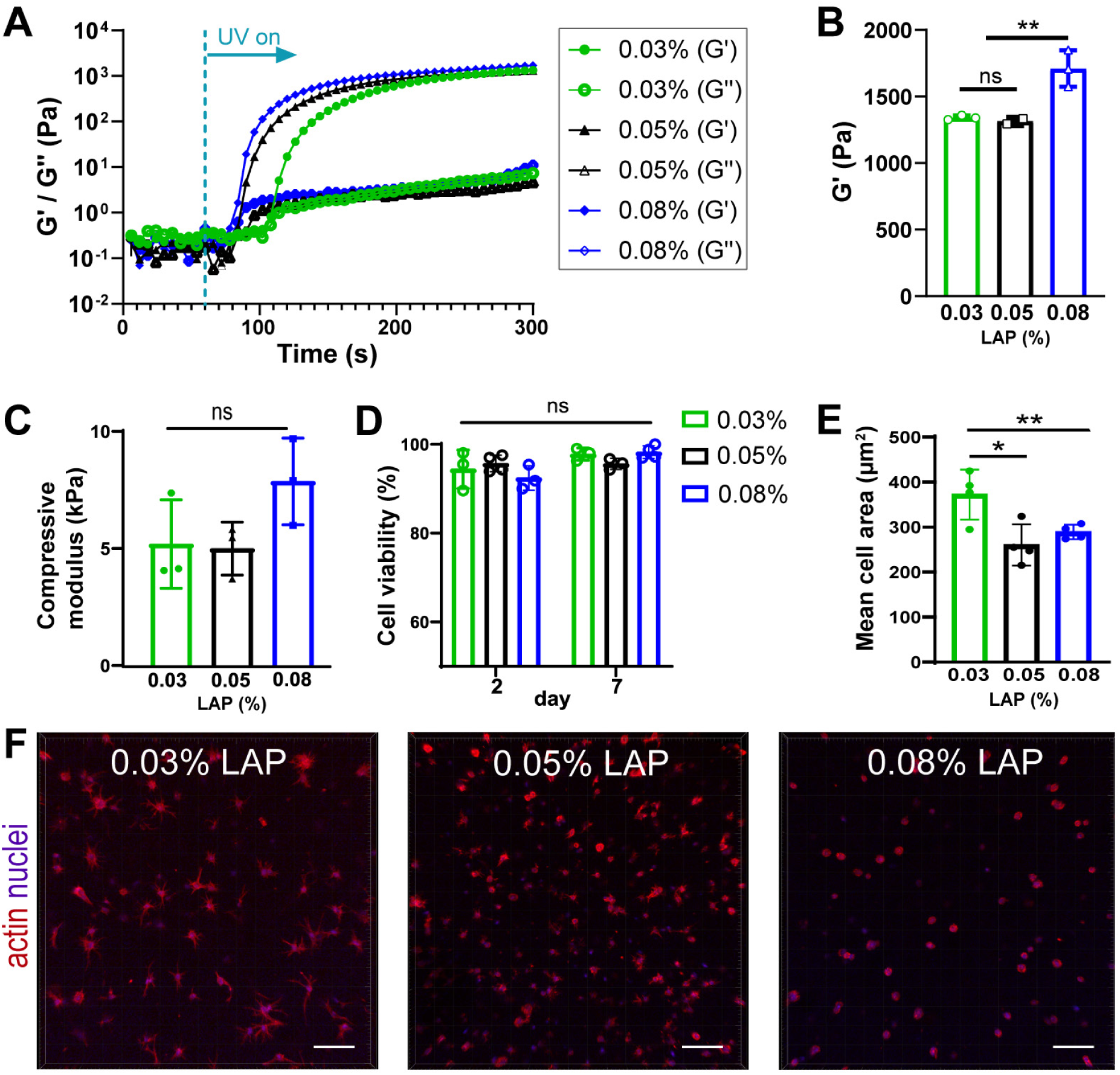
Effects of LAP photoinitiator concentration on gel crosslinking, stiffness, and cell activity. All experiments were performed with resins containing 5% GelMA and different LAP concentrations (0.03%, green; 0.05%, black; 0.08%, blue) **A)** Time sweep of storage and loss moduli (G’/G’’) of acellular resins with UV-365nm light after 60s (dashed line) (n=3). Data points represent the mean, error bars are not shown for better visibility. **B)** Effect of LAP concentration on G’-plateau values determined by photo-rheology. **C)** Mechanical characterization of acellular printed gel constructs after incubation in PBS for 24h (n=3). **D)** Viability of cells in printed constructs on day 2 and 7 (n=3-4). **E)** Average surface area of embedded cells on day 7 as indicator for cell spreading and dendrite formation. **F)** Cell morphology in printed constructs at day 7, visualized by actin-nuclei staining, maximum intensity projection (MIP), scale bars =100 µm. Columns and error bars represent mean and standard deviation (B-E), symbols represent data points; ns ≙ p > 0.05, *p < 0.05, **p < 0.01, ***p < 0.001.

Additionally, the compressive moduli of 3D printed acellular GelMA constructs after swelling was determined with a Zwick mechanical testing machine (**Figure 4C**). Stiffness differences between the scaffolds with the two lower (0.03% and 0.05% LAP ≙ 5 kPa) and the one higher (0.08% LAP ≙ 8 kPa) LAP concentration were observed, although not being statistically significant. These results are consistent with the previously reported observations of faster crosslinking due to higher number of free radicals [29].

Next, we varied LAP concentrations to investigate the influence of LAP and the respective required light dose on cell-compatibility and permissiveness of printed hydrogel environments with cell viability and morphology assays. A high cell viability of >92% after 7 days was reached for all constructs without differences (**Figure 4D, Figure S4**). However, remarkable differences were observed when assessing the morphology of embedded cells after an actin-nuclei staining. Cell spreading with fine dendritic processes could be observed after 7 days for the samples with 0.03% and 0.05% LAP concentration (**Figure 4F**; **Supplementary Video S2**), whereas cells remained round in 0.08% LAP containing gels. This suggests that higher LAP concentrations, and the thereby increased exposure of cells to free radicals might pose toxic effects to the cell fidelity without affecting viability directly [35, 38, 39]. Although previous studies have evidenced negligible cytotoxic effects of soluble GelMA [32] and LAP [35], a thorough assessment of possible phototoxicity mechanisms such as reactive oxygen species[33] during UV irradiation is warranted in future investigations. The observation of increased cell spreading was confirmed by an increased cell surface area in constructs with lower LAP concentration (**Figure 4E**). These findings explain the observed delayed osteogenic differentiation of fully embedded cells compared to top-seeded cells on acellular constructs or 2D cultures [9, 18]. Therefore, 5% GelMA with 0.05% LAP was identified as bioresin of choice with optimal handling and permissiveness and was used for the following cell culture and functional analysis studies.

### 3.2 Enhancing hMSC osteogenesis with endothelial co-cultures

Next, we evaluated whether endothelial co-cultures enhanced *in vitro* osteogenic differentiation and functionality in the 3D printed constructs. Therefore, we assessed marker gene expression by qPCR, performed enzyme activity assays and followed the level of matrix secretion by mechanical testing. A co-culture system was established using stem cells in the osteogenic lineage and endothelial cells (HUVEC), which have the capacity to self-organize and establish intercellular connections. The suitability of the used co-culture medium for both cell types was confirmed by a high viability of >97% and unaltered morphology after 7 days (**Figure S6**).

Relative gene expression of the osteoblastic and osteocytic markers (*ALPL, RUNX2, BGLAP, COL1a2, PDPN, DMP1, SOST*) in mono-and co-cultures (i.e., hMSC and hMSC/HUVEC) were quantified by qPCR weekly (**Figure 5**; **Table S1**). Mono-cultures of HUVECs were used as the control. Importantly, neither considerable amounts of ALP enzyme activity, nor reasonable expression of any of the relevant osteogenic genes could be observed in HUVEC mono-cultures. Therefore, the contribution of HUVECs to total osteogenic gene expression levels in co-cultures was neglected (**Figure S7B**). Generally, all osteoblastic marker genes were expressed throughout the cultivation time, indicating ongoing osteogenic differentiation.

**Figure 5:**
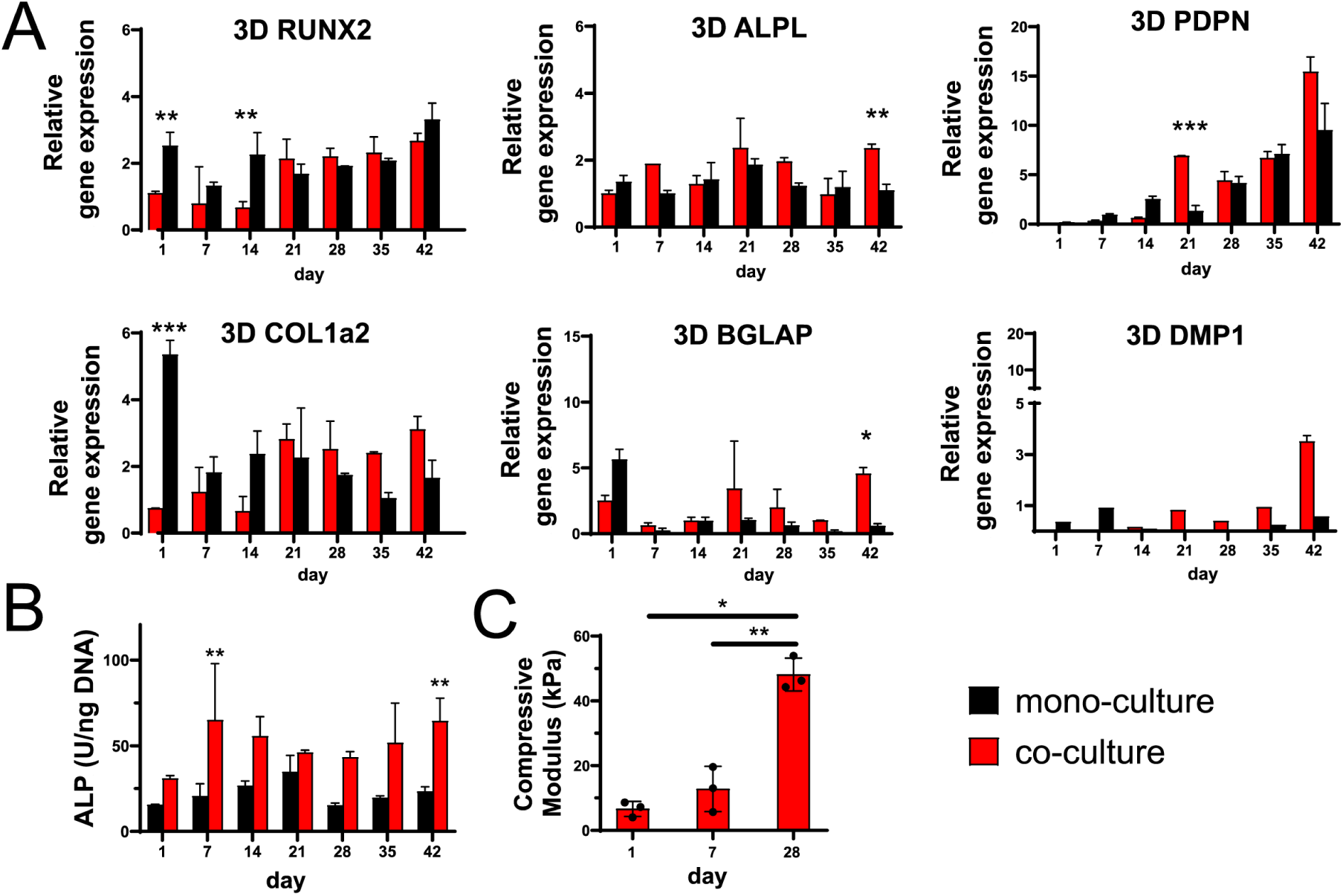
Functional analysis of osteogenic differentiation in co- and mono-cultures in 3D. **A)** Gene expression analysis of osteoblastic (RUNX2, COL1a2, ALPL, BGLAP) and osteocytic (PDPN, DMP1) marker genes over 6 weeks (n= 2-3). Graphs compare mono-with co-cultures in 3D printed cellular constructs. **B)** ALP activity in osteogenic cultures was normalized to the DNA content (n=2-3). **C)** Compressive moduli of co-cultured samples after 1, 7 and 28 days of cultivation (n=3). Columns and error bars represent mean and standard deviation, symbols represent data points (D); *p < 0.05, **p < 0.01, ***p < 0.001.

At the early stage of osteogenesis, hMSCs showed a higher expression of several osteogenic markers in the mono-cultures. The transcription factor *RUNX2* is a key marker for the pre-osteoblast to osteoblast transition. As shown in **Figure 5A**, *RUNX2* expression appeared to be significantly higher in mono-cultures than co-cultures at the early osteogenic stage on day 1 and 14. From 21 days on, its expression in co-culture reached a similar level to mono-culture. A similar trend was observed in the early-stage expression of *COL1a2* genes, indicating the secretion of Collagen-I as a matrix protein by hMSC-derived bone cells. After 21 days, an increase of *COL1a2* expression in the co-cultures compared to mono-cultures was observed.

The *ALPL* encodes for ALP which is synthesized and secreted by osteoblasts for matrix mineralization. On day 42 a significantly higher expression of *ALPL* was found in co-cultures.

The gene *BGLAP* codes for osteocalcin, a bone derived hormone that is solely secreted by osteoblasts. Although on day 1 it was higher expressed in mono-cultures, after 21 days the level of expression significantly increased in the co-cultures (**Figure 5A**). MMP14, also known as MT1-MMP, is a proteolytic enzyme involved in matrix remodeling during osteogenesis and is predominantly active in mature osteoblasts [40]. It was continuously highly expressed and no significant differences between co- and mono-cultures could be observed (**Figure S7A**). Together, at the early stage of 3D osteogenesis most osteoblastic marker genes showed a higher level of expression in mono-cultures and then a trend of enhanced gene expression in co-cultures at later timepoints was identified. These findings imply the temporal dynamics of gene expression in 3D osteogenesis as well as the significant role of juxta- and/or paracrine signaling in cell activity.

In accordance with this trend, marker genes of early osteocytes were increasingly expressed in 3D co-cultures. The early osteocytic marker gene *PDPN* codes for a transmembrane glycoprotein which is important for dendrite elongation. After day 21, it was consistently higher expressed in 3D co-cultures compared to mono-cultures with the highest reached expression levels after 42 days. Moreover, the early osteocytic marker DMP1, a regulator for matrix mineralization, was higher or exclusively expressed in co-cultures compared to mono-cultures, especially in a later stage after day 21. No expression of the marker gene *SOST* for mature osteocytes was observed at any time point. Similar to the effects observed on the expression of osteoblast markers, co-cultures tend to the enhance expression of early osteocytic marker genes.

To confirm the *ALPL* gene expression, an ALP activity assay was performed (**Figure 5B, Figure S8B**). HUVEC controls showed no considerable ALP activity which allowed direct comparison between 2D/3D mono-and co-cultures (**Figure S7A**). ALP activity in 3D was not matched to the *ALPL* gene expression patterns and rather stable expressed in both, mono-and co-culture, during the whole observation time. ALP activity in co-cultures was generally increased compared to mono-cultures with significant differences on day 7 and day 42.

The differences in gene expression and enzyme activity between mono- and co-cultures we also monitored in 2D cultures, additionally to the characterization of 3D printed constructs (**Figure S8**). Interestingly, we observed the effects of co-cultures on osteogenic differentiation being less prominent in 2D cultures compared to 3D. This observation can be explained by the potential of 3D environments to allow endothelial self-organization into vasculature-like structures, and related alterations in signaling [41]. However, further validation by cell tracing, cell tracking and molecular analysis is required to prove this hypothesis. Presence of a self-organized vasculature-like network would promote cell-cell communication, nutrient transport and tissue maturation for a long-term functional 3D culture, even though top-seeding approaches onto porous scaffolds are still frequently used [9]. Additionally, it has to be noted that a direct comparison of 2D and 3D cultures in this study can only be done with a certain reluctance since cell number, spatial organization, culturing environment and RNA isolation techniques differed significantly.

Gene expression analysis and enzymatic assays are powerful tools to assess construct functionality. But especially for bone tissue formation, the level of matrix mineralization is an important indicator. To evaluate this, we followed matrix mineral deposition by time-lapsed micro-CT scans. Printed mono- and co-cultures were cultivated in bioreactors and scanned weekly for 6 weeks, but no sufficient mineralization could be visualized at any time point. Thus, we investigated whether other matrix parameters such as the gel mechanics changed over time. Changes in the compressive moduli of 3D co-culture samples were monitored by mechanical testing on day 1, 7 and 28 after printing. Here, the compressive moduli of the gel constructs increased from about 6 kPa (d1) to 13 kPa (d7) and 46 kPa (d28), with an 8-fold increase between day 1 and 28 (**Figure 5C**). Moreover, an increasing level of gel turbidity was observed during 3D co-culture (**Figure S9**). These results imply that there has been remarkable level of matrix secretion during 3D co-culture. Nevertheless, future work is warranted to investigate if there are remarkable differences in mineral formation between co-culture and mono-culture and to devise strategies to promote tissue maturation with a higher level of mineralization.

### 3.3 Volumetric bioprinting of pre-vascularized constructs

Given the promise of our 3D bioprinted co-culture platform, we further explored VBP of a perfusable pre-vascularized bone construct. Using a 5% GelMA bioresin with 0.05% LAP, single hMSCs were printed inside 3D constructs with a hollow channel and pre-differentiated into the osteogenic lineage by cultivation in osteogenic medium (**Figure 6A**). On day 7, bottom-closed channels were loaded with a dense HUVEC cell suspension in a supporting Collagen-I hydrogel, which provides a 3D environment for self-organization of HUVEC cells.

**Figure 6:**
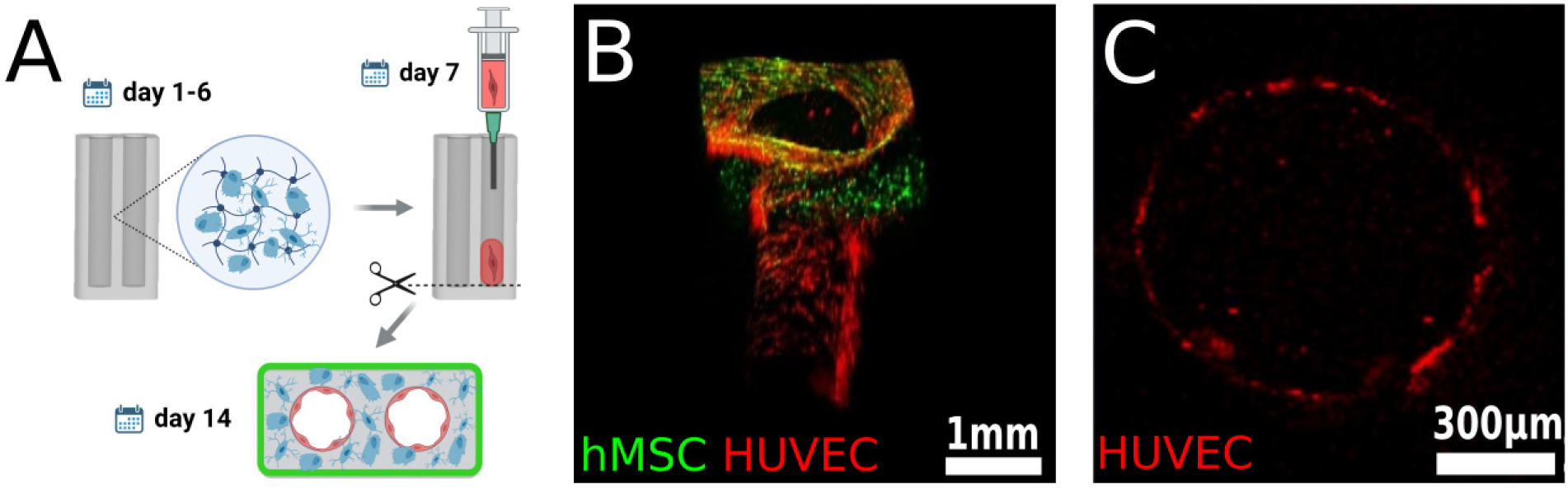
Establishment of a heterocellular perfusable pre-vascularization model. **A)** Schematic experimental procedure of endothelial channel lining in osteogenic 3D constructs: pre-differentiation of hMSC-containing gel constructs for 7 days, injection of endothelial cells with a supporting collagen matrix into the channels, removal of hydrogel plugs to restore medium perfusion, self-organization of endothelial cells into channel lining after 14 days. Created with BioRender.com. **B)** 3D rendered confocal image of an endothelium-lined channel on day 14. hMSCs on the construct surface were stained with calcein-AM (green), while HUVECs inside the channel were stained with DiD (red), respectively. Scale bar = 1 mm. **C)** Cross-section confocal image of an endothelium-lined channel on day 14. HUVECs stained with DiD (red). Scale bar = 300 µm.

After initial cell seeding, perfusion of channels was restored by removal of covering hydrogel plugs. Confocal 3D imaging after additional 7 days of cultivation confirmed the lining of channels with a self-organized endothelial monolayer (red) (**Figure 6B-C**; **Supplementary Video S3**). This pre-vascularized channel model could also be used together with millifluidic perfusion systems to study how flow-induced shear stress promotes cell maturation and matrix mineralization [42]. Our prototype paves the way for adaptor-free perfusion and can eventually drive the development of personalized transplants. However, the resolution of VBP and the endothelial cell seeding methods have to be optimized to reach physiological microvascularization [43].

## 4. Conclusion & Outlook

In conclusion, we demonstrated ultrafast tomographic VBP of cm-scale cell-laden hydrogel constructs for enhanced *in vitro* osteogenesis through 3D endothelial co-culture. A systematic screening of bioresin compositions such as polymer and initiator concentration identified a soft bioresin (5% GelMA, 0.05% LAP) that combines good printability and cell-compatibility for tomographic photopatterning. Compared to the benchmark bioresin of 10% GelMA, our bioresin is much softer and offers enhanced permissiveness for cell-matrix remodeling and cell-cell communication in 3D. Moreover, we performed functional analysis of hMSC/HUVEC 3D co-cultures and hMSC mono-cultures for six weeks. The results revealed enhanced expression of early osteocytic markers (PDPN and DMP1) in the 3D bioprinted constructs under co-culture after 21 days, implying the accelerated osteogenic differentiation by up-regulated juxta- and paracrine signaling in the co-culture system. Additionally, a perfusable pre-vascularized construct with endothelial cell lining was successfully established. To the best of our knowledge, this is the first study to rapidly fabricate perfusable bone-like constructs by VBP.

Nevertheless, the limited level of marix mineralization in combination with the absence of mature osteocytic gene signatures (SOST) emphasize the need to further improve our strategies for *in vitro* osteocytic differentiation. Implementation of the optical tuning method with Iodixanol [27] may enable VBP of bone-like constructs at higher cell densities to accelerate tissue maturation. Other studies have identified additional parameters that can further drive osteogenesis, such as co-culture with macrophage/osteoclast [44] and mechanical stimulation [45] which is a crucial cue for matrix mineralization. The described *in vitro* platform can be upgraded with mechanical loading experiments to enhance *in vitro* osteogenesis towards higher level of functionality by employing spinning or dynamic compression bioreactors [9, 46]. In future, we envisage that researchers can take advantage of this co-culture VBP platform for scaled fabrication of more complex human tissues within seconds for applications in regenerative medicine and *in vitro* drug discovery.

## Acknowledgements

The authors would like to thank Leana Bissig for experimental support with bio-imaging and Christian Gehre for assistance with qPCR. J.G. acknowledges the financial support from the Heyning-Roelli Foundation. W. Qiu gratefully acknowledges the financial support from the China Scholarship Council (CSC). X.H.Q. is grateful to the financial support by the Swiss National Science Foundation (Project No. 190345).

## Author Contributions

J.G. and X.H.Q. conceived the study and designed the experiments, J.G. performed most of the experiments including 3D bioprinting, mono- and co-cultures, confocal laser microscopy imaging and molecular biology assays. W.Q. performed mechanical testing of the hydrogels and analyzed the data. G.N.S. performed micro-CT imaging and data analysis. All authors analyzed the results and participated in the manuscript writing, X.H.Q. and R.M. jointly supervised this research.

## Notes

### Competing Interest Statement

The authors have declared no competing interest.

### Summary of Updates

- Graphical Abstract - Figure 1 - Figure 2 (microCT structural analysis) - Figure 3 (schematic optimization) - Conclusion

