## Supplementary Information for "Tomographic Volumetric Bioprinting of Heterocellular Bone-like Tissues in Seconds"

**Supplementary Figures**

**Figure S1**

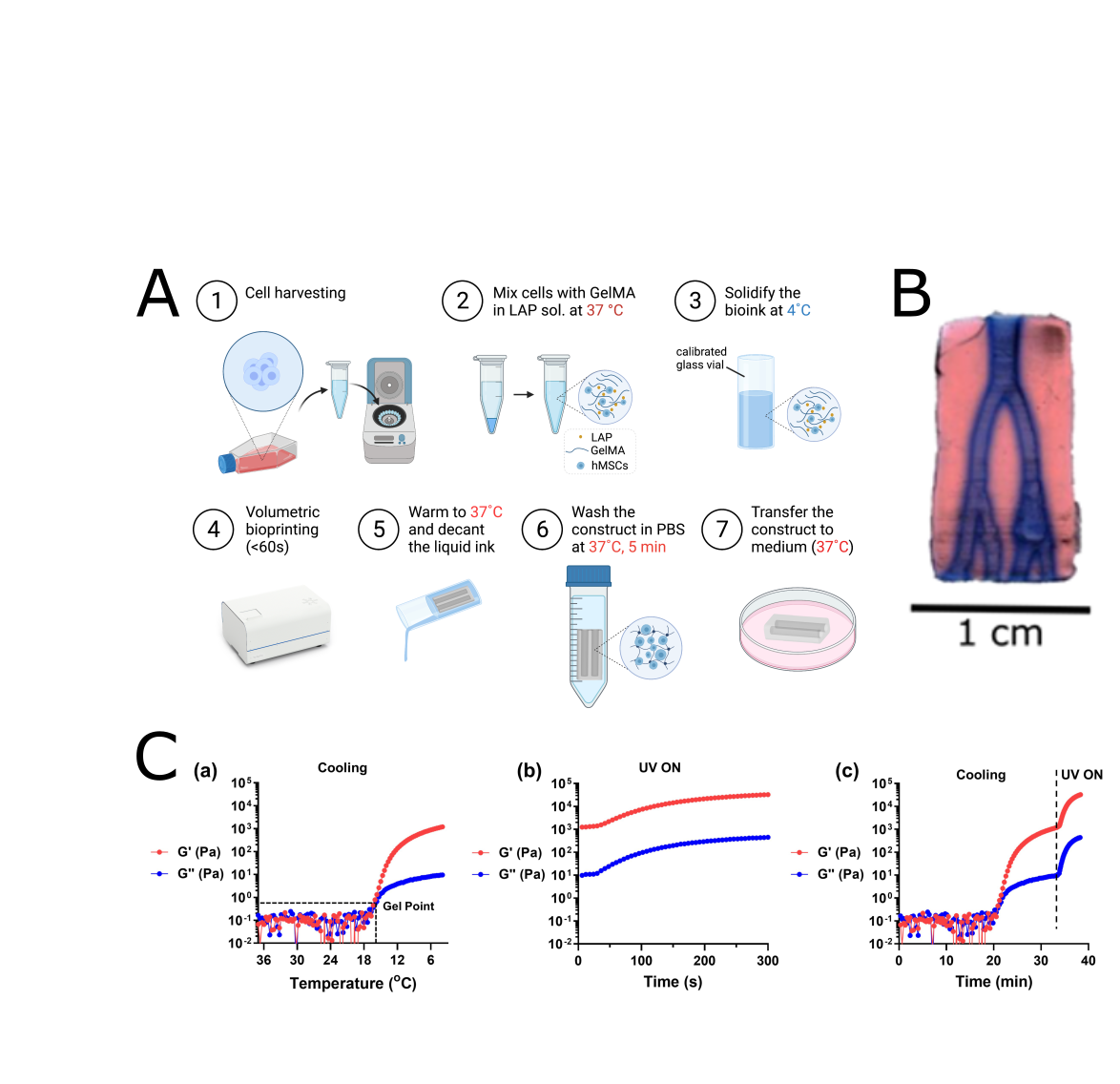

**Figure S1: Volumetric bioprinting process and photorheological characterization of resin formulations.** **A)** Schematic of the sample preparation, volumetric printing process, and construct cultivation. Created with BioRender.com. **B)** Photograph of a cm-scale printed vascular branch model with initial EosinY (pink) staining and subsequent perfusion of channels with Alzian blue (blue). **C)** To mimic the volumetric printing process, a multi-step rheology measurement of 5% GelMA with 0.05% LAP was performed. (a) Storage (G’) and loss modulus (G’’) during an oscillatory time sweep while cooling from 37°C to 4°C at a rate of 1 °C/min. Sol-gel transition at 16˚C. (b) Subsequent exposure to UV-LED 365 nm light (10 mW/cm^-2^) for covalent crosslinking of GelMA via free radical polymerization resulted in a rapid increase in shear modulus (G’ ~ 3.2 kPa, G’’ ~ 450 Pa) after 5 min irradiation. (c) Combined rheology plots that recapitulates the printing process (c). Strain, 1%; frequency, 1 Hz.

**Figure S2**

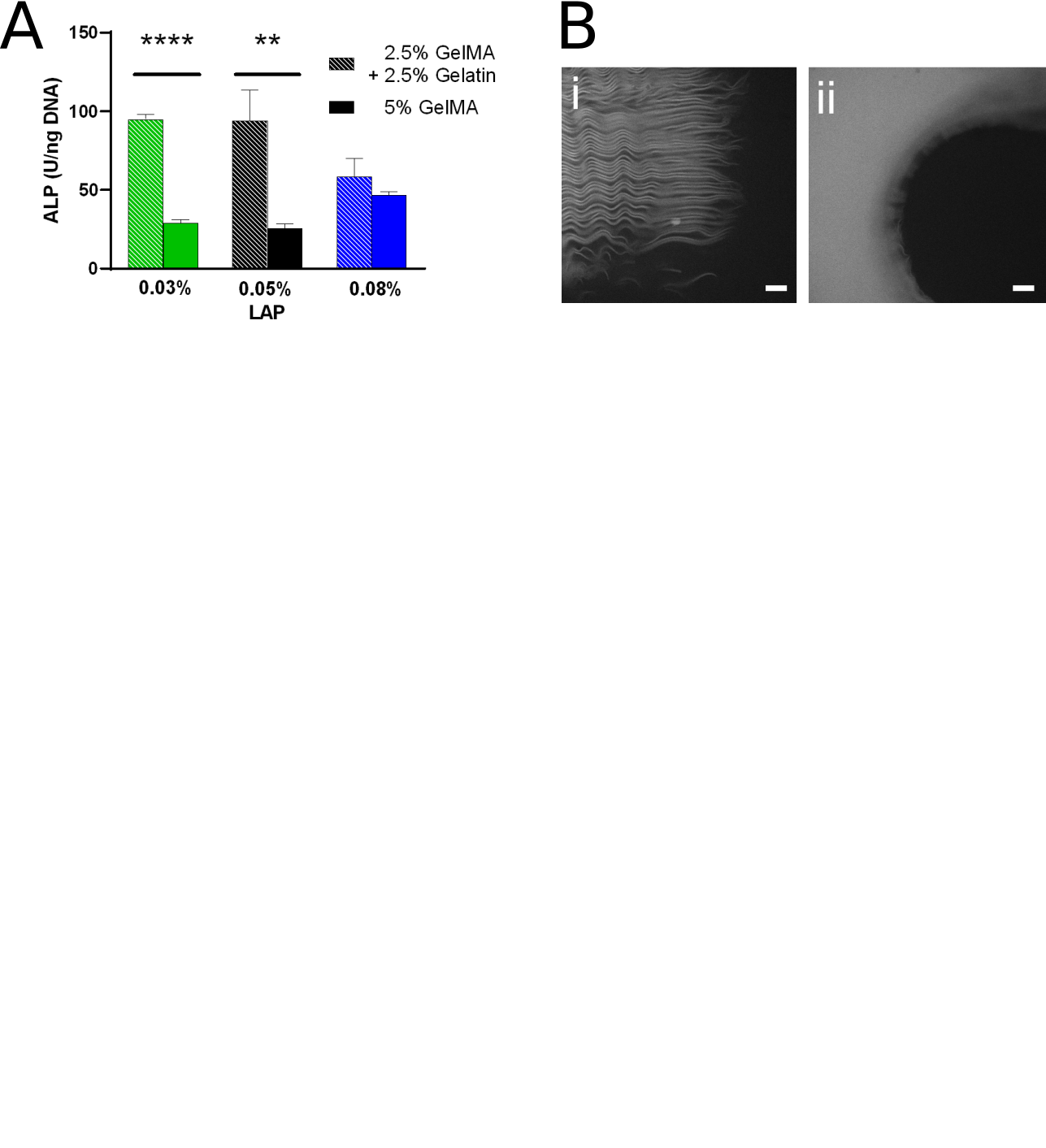

**Figure S2: Characterization of GelMA constructs with sacrificial gelatin**. Volumetric printing can be utilized for the fabrication of 4D constructs with time-dependent changes in stiffness and porosity. Therefore, the printing resin GelMA can be partially substituted with unmodified gelatin which will gradually diffuse out with ongoing incubation at 37°C. **A)** ALP expression of 3D hMSC mono-cultures after 21 days of cultivation in osteogenic medium, normalized to the total DNA content. Comparison of constructs fabricated with either only 5% GelMA or 5% GelMA/gelatin mixture (2.5% GelMA + 2.5% gelatin), and different LAP concentrations (0.03-0.08%). **B)** Confocal fluorescence microscopy images after EosinY staining of printed constructs (3 days after print) revealing the decreased surface smoothness of constructs with sacrificial gelatin (2.5% GelMA, 2.5% gelatin with 0.05% LAP) (side view: i; top view: ii), scale bars 100 µm.

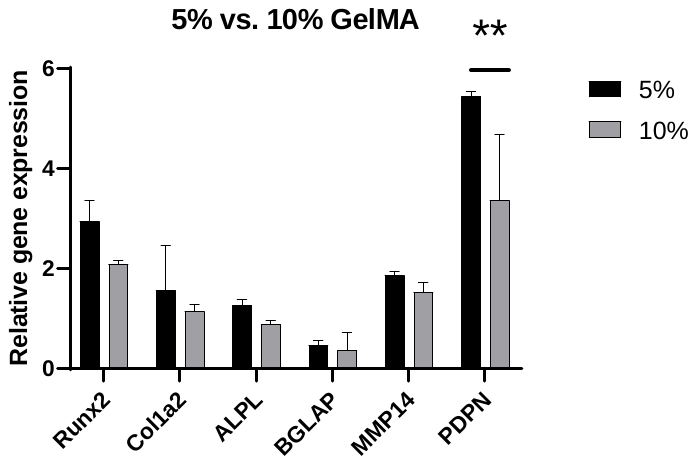
**Figure S3**

**Figure S3: Comparison of relative osteogenic gene expression in 3D constructs containing 5% and 10% GelMA.** Osteogenic 3D mono-cultures were cultivated for 46 days and the relative gene expression of selected marker genes was determined (n=3). Columns and error bars represent mean and standard deviation; **p < 0.01; t-tests with Holm-Sidak correction were used to highlight differences between constructs with 5% and 10% GelMA for each gene. Increased expression of osteoblastic and osteogenic genes (esp. PDPN) was observed for constructs with lower GelMA concentration.

**Figure S4**

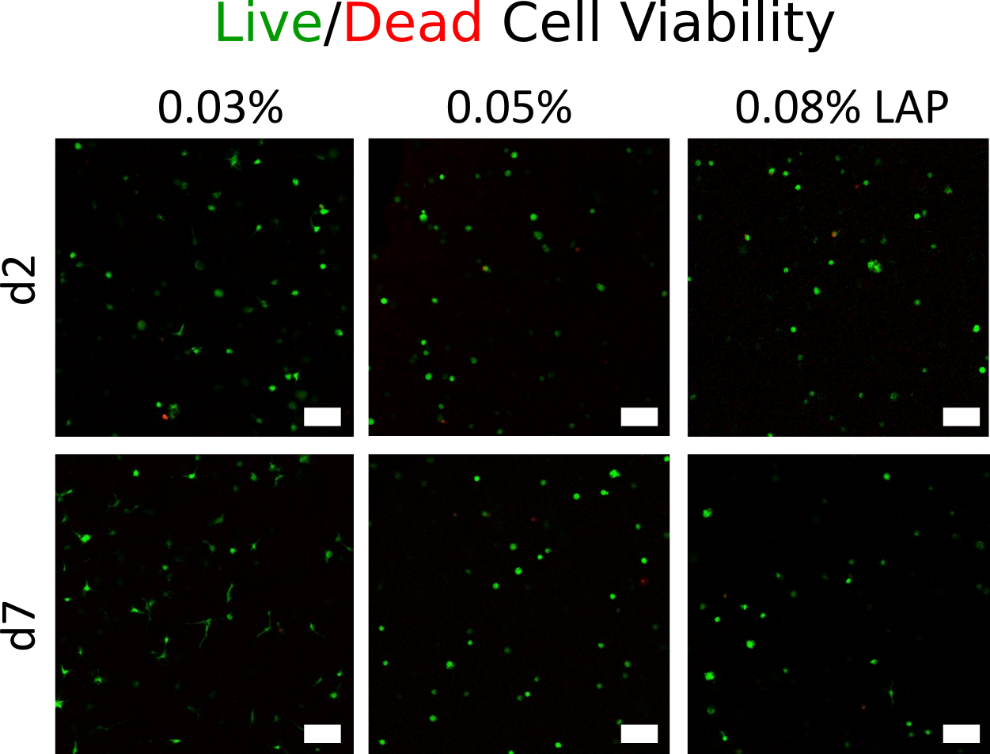

**Figure S4: Viability of cells inside printed constructs on day 2 and 7.** Confocal fluorescence microscopy images of cellular constructs after life-dead staining with Calcein-AM (green, life) and Ethidium-homodimer (red, dead). Cell numbers from the images (n=3-4) were calculated using the automated particle analysis plugin on ImageJ (NIH). Scale bars = 20µm.

**Figure S5**

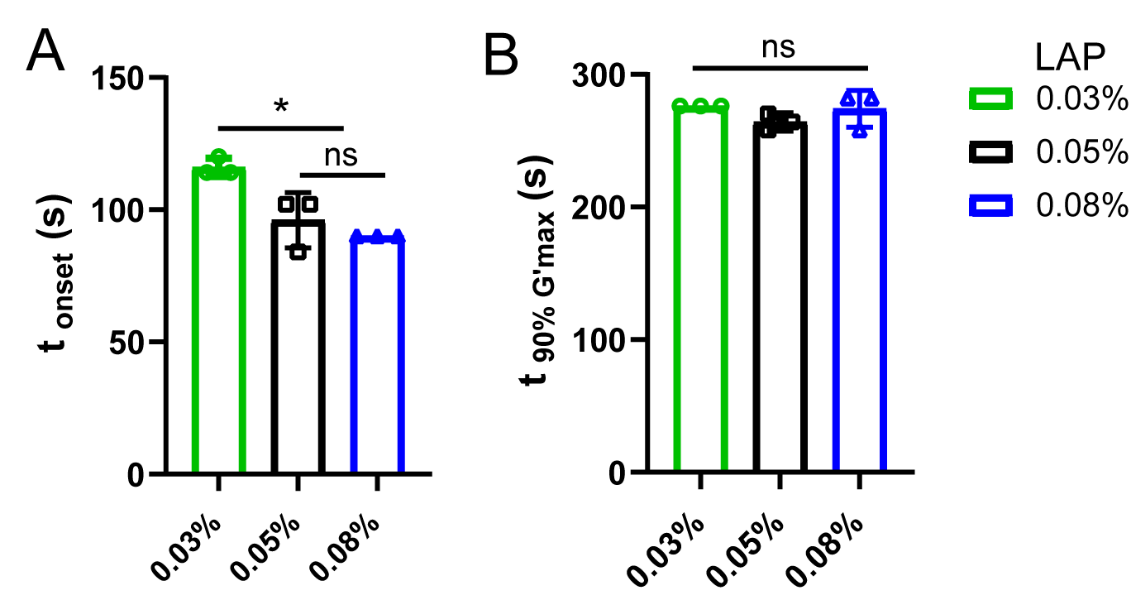

**Figure S5: Effect of LAP initiator concentration on the photocrosslinking of 5% GelMA determined by *in situ* photo-rheology: t_onset_ (A) and t_90% G’max_ (B).** ‘‘*’’ denotes p < 0.05, whereas ‘‘ns’’ denotes no significant difference (p > 0.05). Data presented as mean ± SD. One-way ANOVA was used for statistical analysis.

**Figure S6**

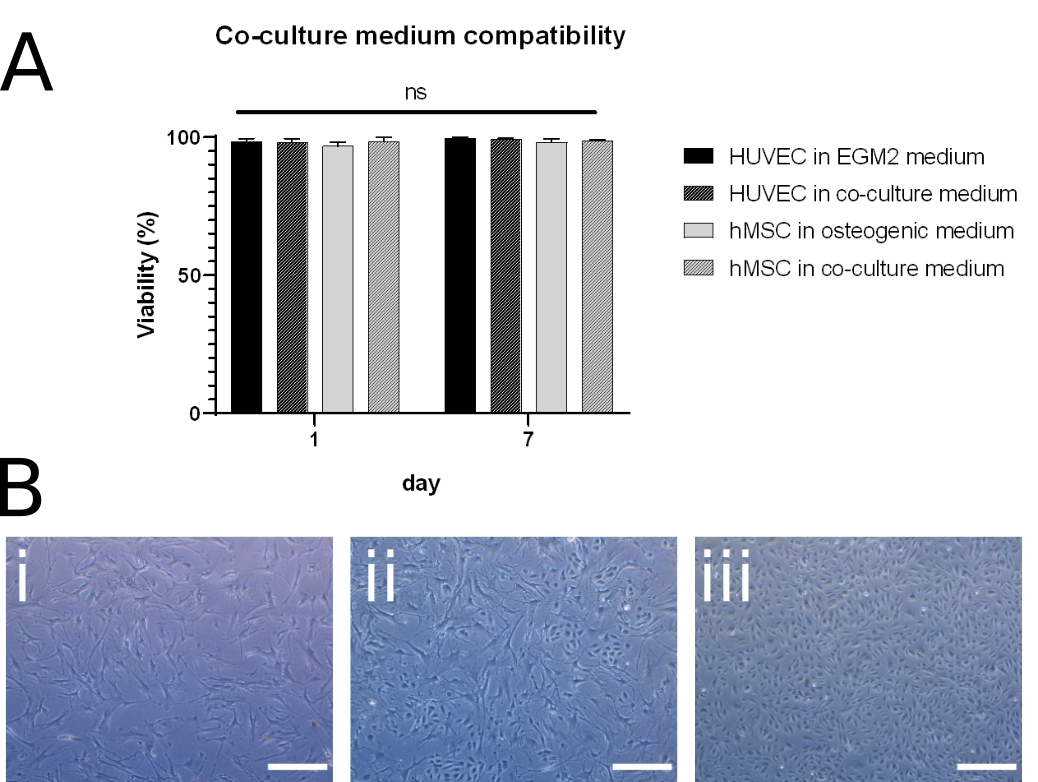

**Figure S6: Co-culture establishment.** Co-cultures of endothelial cells (HUVEC) and mesenchymal stem cells (hMSC) were established with the aim to accelerate osteogenic differentiation. Therefore, a ratio of 1:5 for HUVEC to hMSC was chosen and co-cultures were cultivated in medium mixed from osteogenic medium and EGM-2 HUVEC medium (1:1). **A)** Confirmation of co-culture medium compatibility with 2D cultures of both cell types (HUVEC and hMSC) after 1 and 7 days of cultivation. Viability was determined after calcein/ethidium homodimer staining (n=2). **B)** Morphology of hMSC mono-cultures (i), hMSC-HUVEC co-cultures (ii) and HUVEC mono-cultures (iii) after cultivation for 7 days in the respective mono-culture medium (i,iii) or in co-culture medium (ii). Phase contrast images were taken with 10x magnification, inverted microscope Zeiss, scale bars = 1 mm.

**Figure S7**

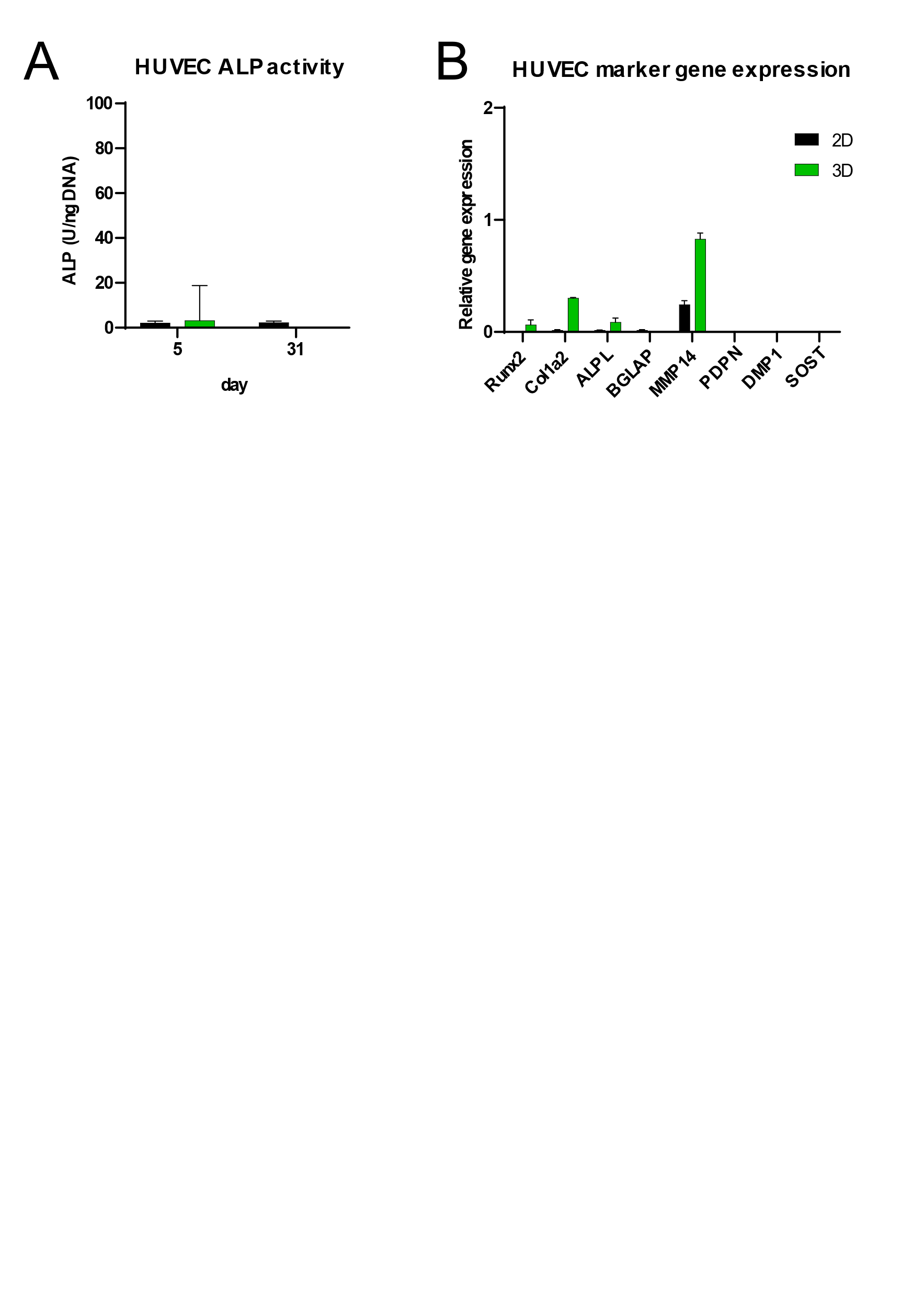

**Figure S7: HUVEC mono-cultures do not express osteogenic markers.** HUVEC mono-cultures were analyzed as negative control for osteogenic characterizations of co-cultures. All constructs were cultivated in co-culture medium. **A)** ALP activity of HUVEC mono-cultures in 2D and 3D was analyzed on day 5 and 31 and normalized to the total DNA content (n=2). **B)** Relative gene expression of osteogenic marker genes in HUVEC mono-cultures after 5 days of cultivation (n=2).

**Figure S8**

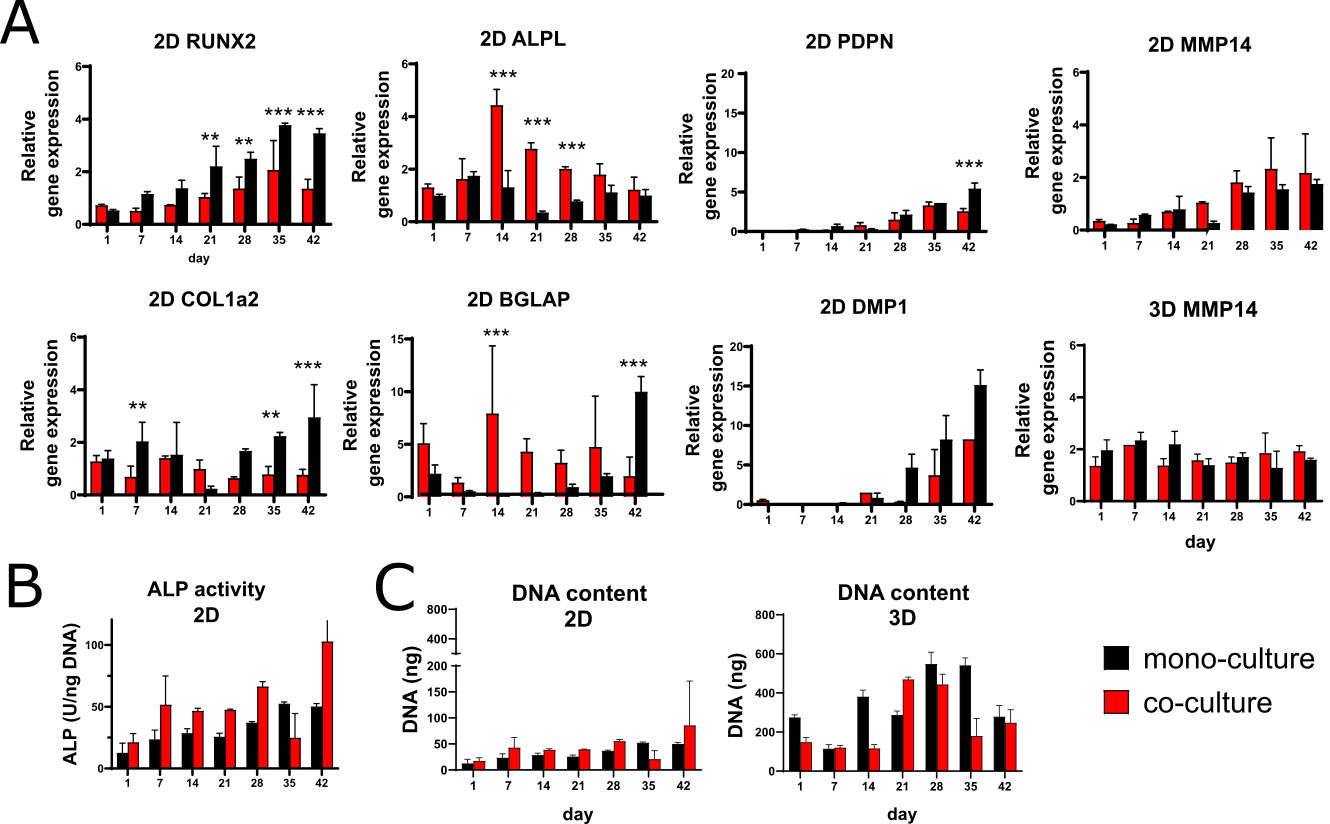

**Figure S8: Functional analysis of osteogenic differentiation in co- and mono-cultures in 2D and 3D (MMP-14). A)** Gene expression patterns of osteoblastic (RUNX2, COL1a2, ALPL, BGLAP, MMP14) and osteocyte-specific (PDPN, DMP1) marker genes to follow osteogenesis over 6 weeks (day 1 - 42) (n=3). Relative gene expression (ΔCt) was normalized to GAPDH and ACTB. Graphs compare mono- with co-cultures in 2D cultures. **B)** ALP enzyme activity in osteogenic cultures was normalized to total DNA content (mono-culture) or estimated hMSC DNA content (co-culture) (n=3). **C)** Total DNA content of osteogenic mono- and co-cultures in 2D and 3D as indicator for cell proliferation (n=3). Columns and error bars represent mean and standard deviation; *p < 0.05, **p < 0.01, ***p < 0.001; t-tests with Holm-Sidak correction were used to highlight differences between co-and mono-culture for each time-point.

**Figure S9**

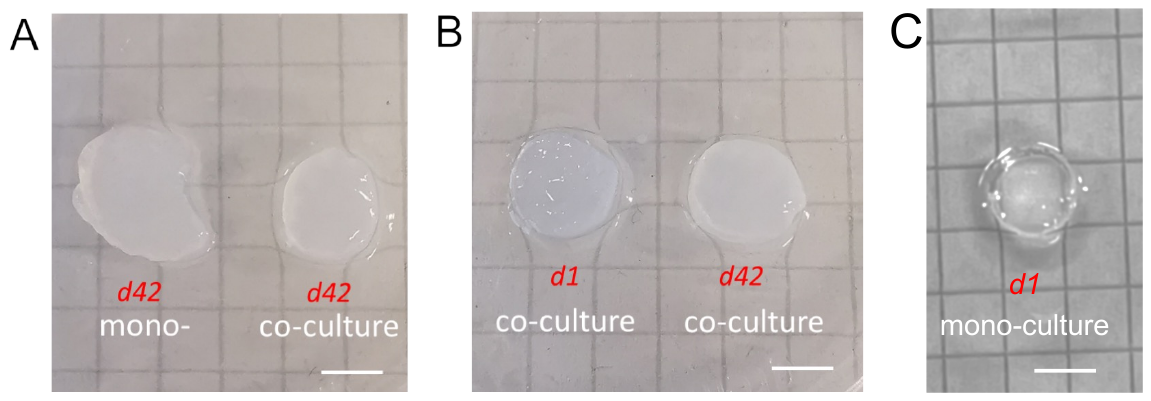

**Figure S9: Photographs of the volumetrically printed samples after mono- and co-culture for 1 day or 42 days.** Turbidity indicates the progress of matrix mineralization. **A)** Left: a gel sample after 42 days mono-culture; right: a gel sample after 42 days co-culture. **B)** Left: a gel sample after 1 day co-culture; right: a gel sample after 42 days co-culture. **C)** Sample after 1 day mono-culture. Scale bars = 5 mm.

**Supplementary Table 1**

**Table S1: Evaluated housekeeping and marker genes for osteocytic differentiation and respective TaqMan probes.**

| **Gene** | **Protein Name** | **TaqMan ID** |
| --- | --- | --- |
| ACTB | Actin Beta | Hs01060665_g1 |
| GAPDH | Glyceraldehyde-3-Phosphate Dehydrogenase | Hs02758991_g1 |
| RUNX2 | Runt-Related Transcription Factor 2 | Hs00231692_m1 |
| COL1A2 | Collagen Type I Alpha 2 Chain | Hs01028956_m1 |
| ALPL | Alkaline Phosphatase, Biomineralization Associated | Hs01029144_m1 |
| BGLAP | Bone Gamma-Carboxyglutamate Protein, Osteocalcin | Hs01587814_g1 |
| MMP14 | Matrix Metalloproteinase 14 | Hs00237119_m1 |
| PDPN | Podoplanin | Hs00366766_m1 |
| DMP1 | Dentin Matrix Acidic Phosphoprotein 1 | Hs01009391_g1 |
| SOST | Sclerostin | Hs00228830_m1 |

**Supplementary Videos**

**Video S1: Volumetric bioprinting procedure**

Printing of a rectangular construct inside a printing tube filled with GelMA 5%, 0.05% LAP. The tomographic light projection can be seen from the left and after several seconds a construct solidifies within the resin. Scale bar, 10 mm.

**Video S2: Cell Spreading in printed GelMA constructs**

3D image of spreading hMSCs with dendrites in 5% GelMA with 0.05% LAP after 7 days of cultivation in osteogenic medium. The video was created with Imaris software.

**Video S3: Endothelial lined channels**

3D image of a volumetrically printed channel with endothelial lining (red) after 14 days of total cultivation and 7 days after HUVEC seeding. hMSCs on the construct surface are stained in green.
